## Supplementary Information for "Cryo-EM structure of coronavirus-HKU1 haemagglutinin esterase reveals architectural changes arising from prolonged circulation in humans"

**Hurdiss et al, 2020.**

### Supplementary Information

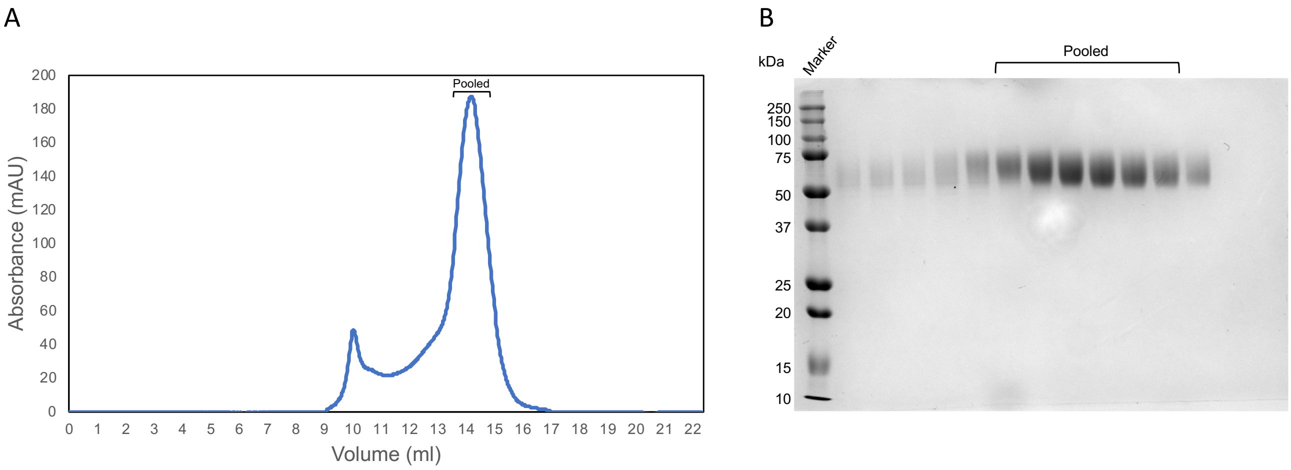


**Figure S1:** Purification of HKU1 HE. A) Elution profile of HKU HE from a Superdex 200 10/300 GL gel filtration column. UV absorbance (mAU) is shown as a blue trace. Fractions pooled for subsequent cryo-EM analysis are indicated. (B) SDS-PAGE analysis of the fractions collected from the gel filtration run shown in (A). The molecular weight marker and pooled fractions are indicated.

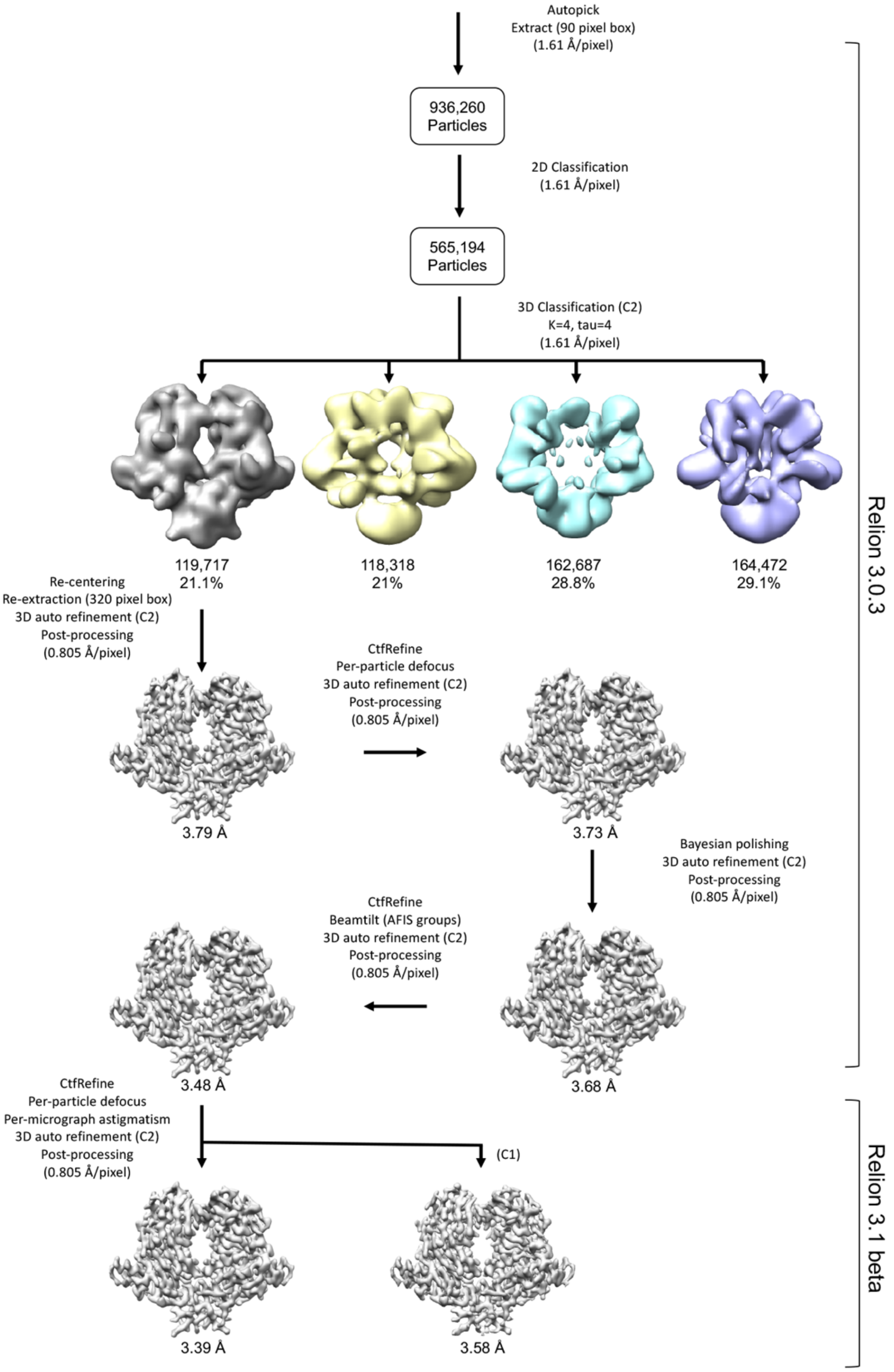


**Figure S2:** Single-particle cryo-EM image processing workflow for HCoV-HKU1 HE (see methods for details).

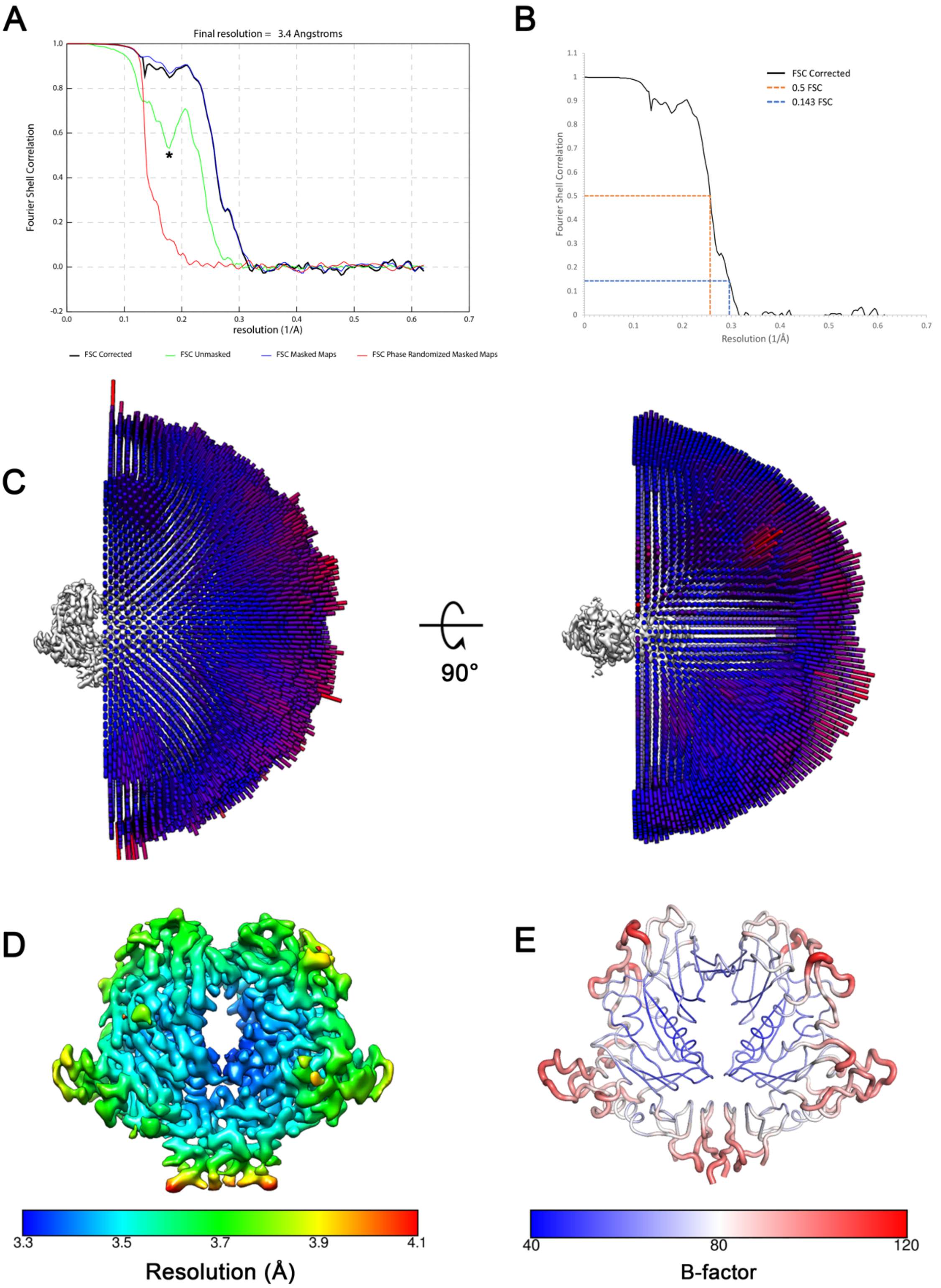


**Figure S3:** A) Gold-standard Fourier shell correlation (FSC) curve generated from the independent half maps contributing to the ~3.4 Å resolution density map. A dip in the unmasked FSC at ~5.8 Å is indicated by an asterisk. This likely results from poor correlation between the disordered and heterogenous glycans at this spatial frequency. B) Corrected FSC curve with the 0.5 and 0.143 cut-off indicated by orange and blue dashed lines, respectively. C) Angular distribution plot of the final C2 refined EM density map. D) The final C2 refined EM density map coloured according to local resolution which was calculated in Relion. E) Atomic model of the dimeric complex with residues coloured according to calculated B-factor.

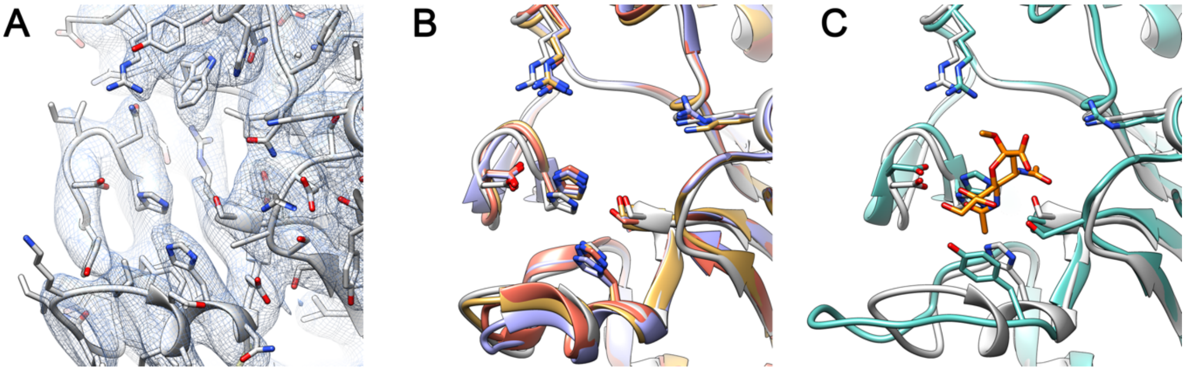


**Figure S4:** A) Zoomed-in view of the HKU1 HE esterase domain active site. EM density is shown as a blue mesh and the atomic model is coloured grey. B) Overlay of the HE esterase domain active-site of HKU1 HE, BCoV (PDB ID:3CL5), MHV-DVIM (PDB ID:5JIF), and OC43 (PDB ID:5N11) coloured grey, purple, gold and red, respectively. Residues which make up the active-site are shown as sticks. C) Overlay of the HE esterase domain active-site of HKU1 HE and MHV-NJ (PDB ID:5JIL) with 4-N-Acetyl sialic acid, present in the latter structure, coloured orange.

```
CLUSTAL O(1.2.4) multiple sequence alignment


HKU1-A              ----------MLIIFLFFYFCYGF-NEPLNVVSHLNHDWFLFGDSRSDCNHINNLKIKNF        49
BCoV-Mebus          ----MFLLLRF-VLVSCIIGSLGFDNPPTNVVSHLNGDWFLFGDSRSDCNHVVNTNPRNY        55
MHV-New-Jersey      --MCIAIAPRTLLLLIGCQLVFGF-NEPLNIVSHLNDDWFLFGDSRSDCTYVENNGHPKL        57
MHV-DVIM            MARTDAMAPRTLLLVLSLGYAFGF-NEPLNVVSHLNDDWFLFGDSRSDCNHINNLSQQNY        59
                                ::.       ** * * *:***** ************.:: *    :

HKU1-A              DYLDIHPSLCNNGKISSSAGDSIFKSFHFTRFYNYTGEGDQIIFYEGVNFNPYHRFKCFP        109
BCoV-Mebus          SYMDLNPALCDSGKISSKAGNSIFRSFHFTDFYNYTGEGQQIIFYEGVNFTPYHAFKCTT        115
MHV-New-Jersey      DWLDLDPKLCNSGRISAKSGNSLFRSFHFTDFYNYSGEGDQVIFYEGVNFSPSHGFKCLA        117
MHV-DVIM            NYMDINPELCKSGKISAKAGNSLFKSFHFTDFYNYTGEGSQIIFYEGVNFTPYVGFKCLN        119
                    .::*:.* **..*:**:.:*:*:*:***** ****:***.*:********.*   ***

                                                         β4-β5
HKU1-A              NGSNDVWLLNKVRFYRALYSNMAFFRYLTFVDIPYNVS--LSKFNSCKS----DILSLNN        163
BCoV-Mebus          SGSNDIWMQNKGLFYTQVYKNMAVYRSLTFVNVPYVYNGSAQSTALCKS----GSLVLNN        171
MHV-New-Jersey      NGDNKIWMGNKARFYARLYEKMAQYRSLSIVTVSYAYGGNAKPTSICKD----NKLTLNN        173
MHV-DVIM            NGDNNRWMGNKARFYTQLYQKMAHYRSLSVINITYTYNGSAGPVSMCKHIANGVTLTLNN        179
                    .*.*. *: **  **  :*.:** :* *:.: : *  .        **       * ***

                           β5-β6                                β7-β8
HKU1-A              PIFIN---------YSKEVYFTLLGCSLYLVPLCLFKSNF--------------SQYYYN        200
BCoV-Mebus          PAYIAREANFGDYYYKVEADFYLSGCDEYIVPLCIFNGKFLS-----NTKYYDDSQYYFN        226
MHV-New-Jersey      PTFISKESNYADYYYVSEANFTLQGCDEFIVPLCVFNGHSRGSSSDPANTYYMDSQMYYN        233
MHV-DVIM            PTFIGKEVSKPDYYYESEANFTLQGCDEFIVPLCVFNGQYLS-----SKLYYDDSQYYYN        234
                    * :*          *  *. * * **. ::****:*:.:              ** *:*

                                   β9-β10
HKU1-A              IDTGSVYGFSNVVY---------PDLDCIYISLKPGSYKVSTTAPFLSLPTKALCFDKSK        251
BCoV-Mebus          KDTGVIYGLNSTETI-----TTGFDFNCHYLVLPSGNYLAISNELLLTVPTKAICLNKRK        281
MHV-New-Jersey      TVTGVFYGFNSTLDVGTTVQNPGLDLTCSYLALSPGNYKAVSLEFLLSLPSKAICLLKPK        293
MHV-DVIM            VDTGVLYGFNSTLNI-----TSGLDLTCIYLALTPGNYISISNELLLTVPSKAICLRKPK        289
                      ** .**:...            *: * *: *  *.*   :   :*::*:**:*: * *

HKU1-A              QFVPVQVVDSRWNNERASDISLSVACQLPYCYFRNSSANYVGK-YDINHGDSGFISILSG        310
BCoV-Mebus          DFTPVQVVDSRWNNARQSDNMTAVACQPPYCYFRNSTTNYVGV-YDINHGDAGFTSILSG        340
MHV-New-Jersey      RFMPVQVVDSRWNSTRQSDNMTAVACQLPYCFFRNTSADYSGDTHDVHHGDLYFRQLLSG        353
MHV-DVIM            AFTPVQVVDSRWHSNRQSDNMTAIACQLPYCYFRNTTSDYNGV-YDSHHGDAGFTSILAG        348
                     * *********:. * **   ::*** ***:***::::* *  :* :***  * .:*:*

HKU1-A              LLYNVSCISYYGVFLYDNFTSIWPYYSFGRCPTSSIIK---HPICVYDFLPIILQGILLC        367
BCoV-Mebus          LLYDSPCFSQQGVFRYDNVSSVWPLYSYGRCPTAADINTPDVPICVYDPLPLILLGILLG        400
MHV-New-Jersey      LLYNVSCIAQQGAFLYNNVSSIWPVYGYGHCPTAANIGY-MAPICLYDPLPVILLGVLLG        412
MHV-DVIM            LMYNVSCLAQQGAFVYNNVSSSWPQYPYGHCPTAANIVF-MAPVCMYDPLPVILLGVLLG        407
                    *:*:  *::  *.* *:*.:* ** * :*:***:: *     *:*:** **:** *:**

HKU1-A              LALLFVVFLLFLLYNDKSH-----    386
BCoV-Mebus          VAVIIIVVLLLYFMVDNGTRLHDA    424
MHV-New-Jersey      IAVLIIVFLILYFMTDSSVRLHEA    436
MHV-DVIM            IAVLIIVFLMFYFMTDSGVRLHEA    431
                    :*::::*.*:: :  *..
```

**Figure S5:** A) Multiple sequence alignment of HKU1-A, BCoV-Mebus, MHV-NJ and MHV-DVIM HE amino acid sequences. The HKU1-A loops deletions are labelled and highlighted yellow.

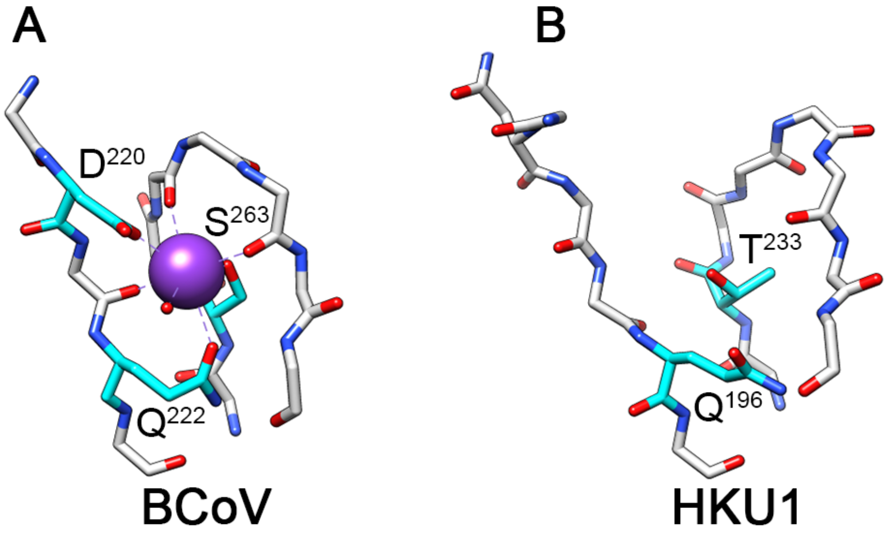


**Figure S6:** A) Stick representation of BCoV's lectin domain metal binding site, with the potassium ion shown as a purple sphere and coordinating sidechains coloured cyan. B) Equivalent view as shown in A for HKU1 HE.

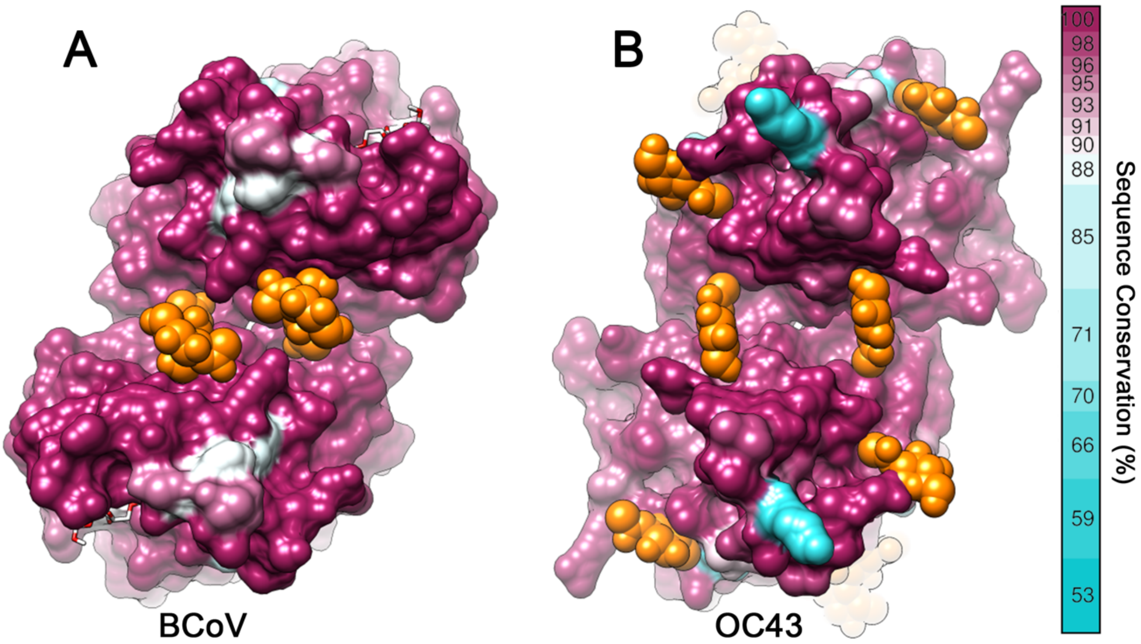


**Figure S7:** A) Top down view of the dimeric BCoV HE lectin domain (PDB ID: 3CL5), shown as a surface representation and coloured according to sequence conservation (75 sequences). B) Equivalent view as shown in A for OC43 HE (65 sequences). The β5-β6 loop of OC43 HE was not modelled in the crystal structure (PDB ID: 5N11) and was therefore modelled, for demonstrative purposes, using equivalent region in BCoV HE.

A

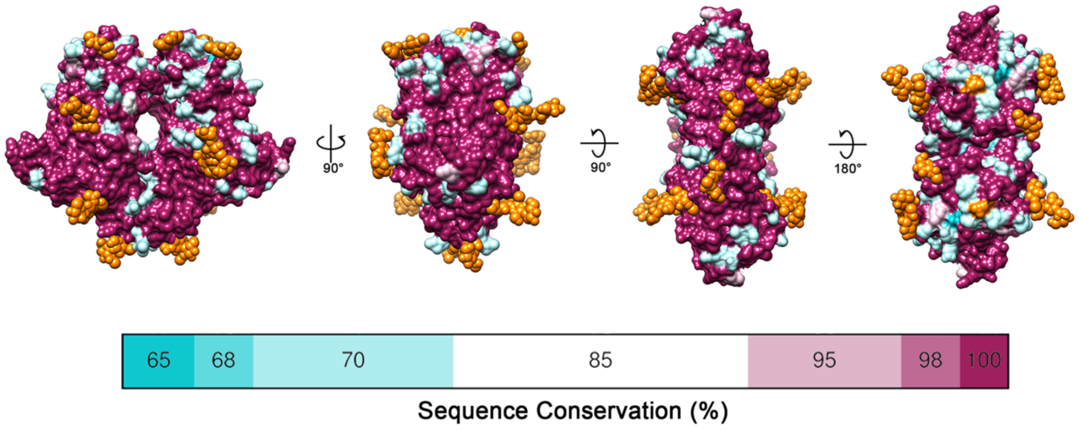


B

```
HKU1-A      MLIIFLFFYFCYGFNEPLNVVSHLNHDWFLFGDSRSDCNHINNLKIKNFDYLDIHPSLCN      60
HKU1-B      MLIIFLFFNFCYGFNEPLNVVSHLNHDWFLFGDSRSDCNHINNLKIKNYGYLDIHPSLCN      60
            ******** ***************************************:.**********

HKU1-A      NGKISSSAGDSIFKSFHFTRFYNYTGEGDQIIFYEGVNFNPYHRFKCFPNGSNDVWLLNK      120
HKU1-B      NGKISSSAGDSIFKSYHFTRFYNYTGEGDQIIFYEGVNFNPHHRFKCSSNGSNDVWIFNK      120
            ***************:*************************:*****  *******::**

HKU1-A      VRFYRALYSNMAFFRYLTFVDIPYNVSLS-KFNSCKSDILSLNNPIFI--NYSKEVYFTL      177
HKU1-B      VRFYRALYSNMALFRYLTFVDILYNFSFSIKANICNSNILSLNNPIFISTNYSKDVYFTL      180
            ************:********* **.*:* * * *:*:**********  ****:*****

HKU1-A      LGCSLYLVPLCLFKSNFSQYYYNIDTGSVYGFSNVVYPDLDCIYISLKPGSYKVSTTAPF      237
HKU1-B      SGCSLYLVPLCLFKSNFSQYYYNMDTGFAYGYSNFVSSDLDCTYISLKPGSYKIFSTGFV      240
             **********************:*** .**:**.*  **** **********: :*. .

HKU1-A      LSLPTKALCFDKSKQFVPVQVVDSRWNNERASDISLSVACQLPYCYFRNSSANYVGKYDI      297
HKU1-B      LSIPTKALCFNKSKQFVPVQVVDSRWNNLRASDTSLSDACQLPYCYFRNSSGNYVGKYDI      300
            **:*******:***************** **** *** *************.********

HKU1-A      NHGDSGFISILSGLLYNVSCISYYGVFLYDNFTSIWPYYSFGRCPTSSIIKHPICVYDFL      357
HKU1-B      NHGDNGFTSILSGLLYNVSCISYYGSFLYDNFTSIWPRFSFGNCPTSAYIK-LNCFYDPL      359
            ****.** ***************** *********** :***.****: **   *.** *

HKU1-A      PIILQGILLCLALLFVVFLLFLLYNDKS        385
HKU1-B      PIILQGILLFLALLFIVFLLFLVYHG--        385
            ********* *****:******:*:.
```

**Figure S8:** A) Orthogonal views of the HKU1 HE model shown as a surface representation, coloured according to sequence conservation (28 HKU1-A sequences and 12 HKU1-B sequences). N-glycans are shown as orange spheres. B) Alignment of representative HKU1-A and HKU1-B HE amino acid sequences. The eight conserved N-linked glycosylation sites are highlighted orange.

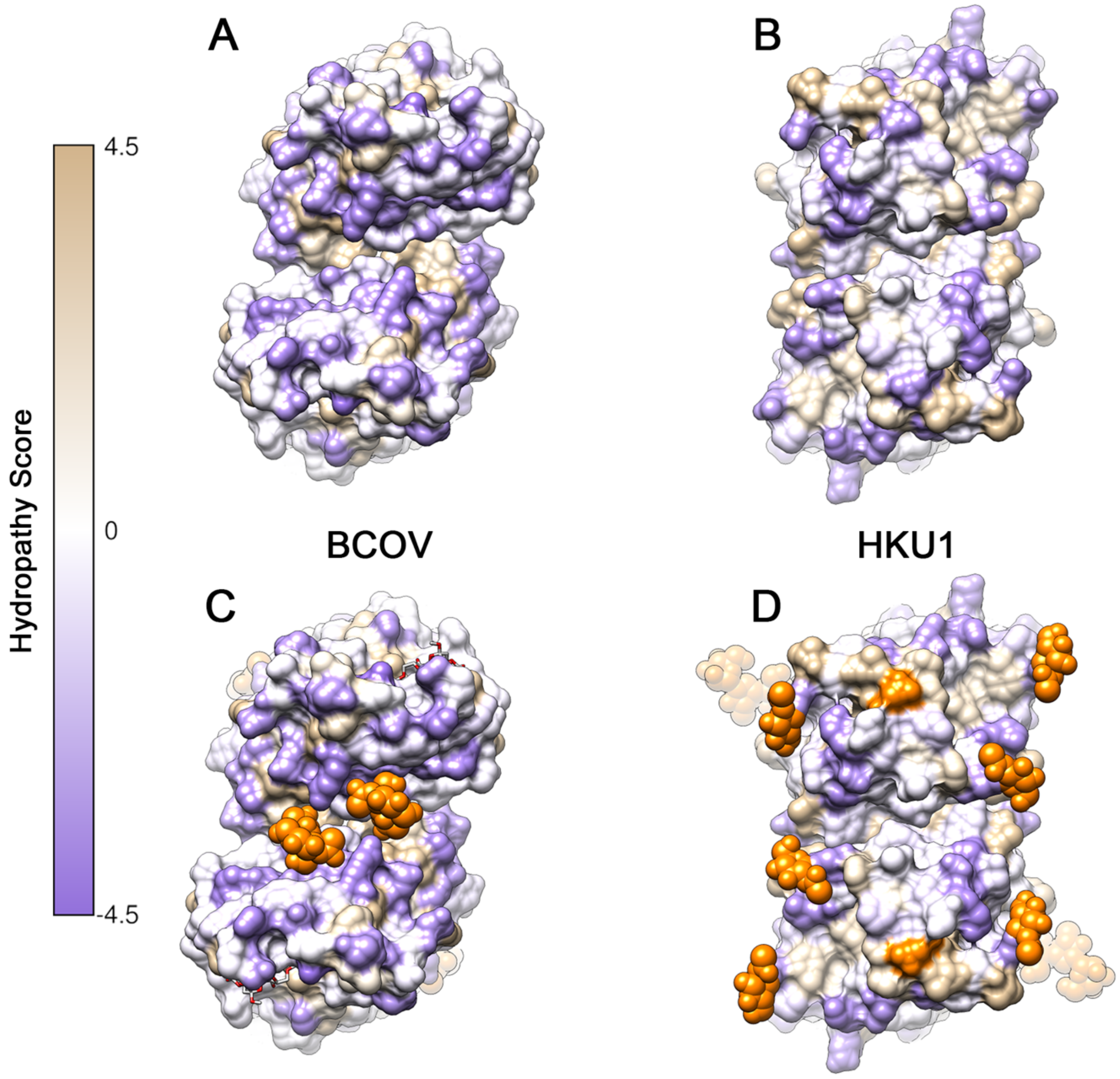


**Figure S9:** A) Top down view of the dimeric BCoV and (B) HCoV-HKU1 HE lectin domains, shown as a surface representation and coloured according to the Kyte-Doolittle scale, where the most hydrophobic residues are coloured tan and the most hydrophilic residues are coloured purple. C-D) As shown in A and B with N-glycosylation sites coloured orange and, if modelled, depicted as spheres. For BCoV, bound sialic acid is shown in stick representation and coloured white.

**Table S1:** Data collection, image processing and refinement information.

| | |
|---|---|
| **Data Collection** | |
| Microscope | Titan Krios G4 |
| Voltage (keV) | 300 keV |
| Nominal magnification | 96,000x |
| Exposure navigation | Image shift |
| Movie acquisition rate | ~238 per hour |
| Detector | Falcon 4 |
| Calibrated pixel size (Å) | 0.805 |
| Cumulative exposure (e/Å$^2$) | 40 |
| Dose rate (e/pixel/sec) | 5 |
| Underfocus range ($\mu$m) | 1 to 2.5 (0.25 increments) |
| Micrographs collected | 6029 |
| **Reconstruction** | |
| Final particles (no.) | 119,717 |
| Symmetry | C2 |
| B-factor (Å$^2$) | -122 |
| **Resolution (Å)** | |
| FSC 0.5 (masked) | 3.89 |
| FSC 0.143 (masked) | 3.39 |
| Resolution range (local) | 3.3-4.8 |
| Angular accuracy | 1.58 |
| **Refinement** | |
| Protein residues/atoms | 664/5414 |
| N-glycans/atoms | 14/440 |
| **Map correlation coefficient** | |
| Mask | 0.8 |
| Volume | 0.79 |
| Peaks | 0.70 |
| **R.M.S. deviations** | |
| Bond Lengths (Å) | 0.007 |
| Bond Angles (°) | 0.972 |
| **MolProbity** | |
| Overall score | 1.79 |
| Clashscore | 4.49 |
| Ramachandran outliers (%) | 0 |
| Ramachandran favoured (%) | 89.55 |
| Rotamer outliers (%) | 0.33 |
| C-beta outliers | 0 |
| **EMRinger Score** | 4.7 |
| **Privateer** | |
| Wrong anomer | 0 |
| Wrong configuration | 0 |
| Unphysical puckering amplitude | 0 |
| In higher-energy conformations | 0 |

**Table S2:** Overview of HKU1-HE glycoproteomics data. Full lists of identified site-specific glycoforms and their semiquantitative analysis are provided per protease dataset.

*The table can be found in the supplementary excel file.*

**Table S3:** Overview of common N-glycoforms shared at the majority of sites with predicted structures.

| Present at No. of glycosites | Predicted components | Predicted structure |
|---|---|---|
| 6 out 8 | HexNac4Hex5Fuc1 | 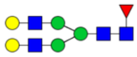 |
| 6 out 8 | HexNac4Hex5Fuc1NeuAc1 | 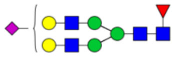 |
| 5 out 8 | HexNac3Hex5Fuc1 | 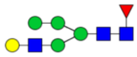 |
| 5 out 8 | HexNac3Hex6 | 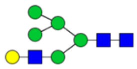 |
| 5 out 8 | HexNac3Hex6Fuc1 | 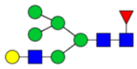 |
| 5 out 8 | HexNac4Hex4Fuc1 | 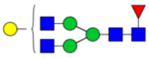 |
| 5 out 8 | HexNac5Hex4Fuc1 | 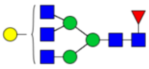 |
| 5 out 8 | HexNac5Hex6Fuc1NeuAc1 | 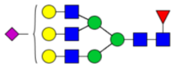 |
| 4 out 8 | HexNac2Hex5 | 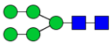 |
| 4 out 8 | HexNac3Hex5 | 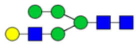 |
| 4 out 8 | HexNac4Hex5 | 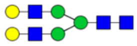 |
| 4 out 8 | HexNac4Hex3Fuc1 | 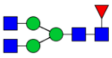 |
| 4 out 8 | HexNac4Hex4Fuc1NeuAc1 | 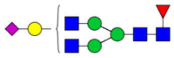 |
| 4 out 8 | HexNac5Hex3Fuc1 | 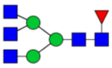 |
| 4 out 8 | HexNac5Hex5Fuc1 | 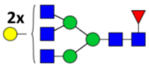  |
